# Pan-Cancer Drug Response Prediction Using Integrative Principal Component Regression

**DOI:** 10.1101/2023.10.03.560366

**Authors:** Qingzhi Liu, Gen Li, Veerabhadran Baladandayuthapani

## Abstract

The pursuit of precision oncology heavily relies on large-scale genomic and pharmacological data garnered from preclinical cancer model systems such as cell lines. While cell lines are instrumental in understanding the interplay between genomic programs and drug response, it well-established that they are not fully representative of patient tumors. Development of integrative methods that can systematically assess the commonalities between patient tumors and cell-lines can help bridge this gap. To this end, we introduce the Integrative Principal Component Regression (iPCR) model which uncovers both joint and model-specific structured variations in the genomic data of cell lines and patient tumors through matrix decompositions. The extracted joint variation is then used to predict patient drug responses based on the pharmacological data from preclinical models. Moreover, the interpretability of our model allows for the identification of key driver genes and pathways associated with the treatment-specific response in patients across multiple cancers. We demonstrate that the outputs of the iPCR model can assist in inferring both model-specific and shared co-expression networks between cell lines and patients. We show that iPCR performs favorably compared to competing approaches in predicting patient drug responses, in both simulation studies and real-world applications, in addition to identifying key genomic drivers of cancer drug responses.

## 1 Introduction

Cancer is a complex disease at multiple levels. Due to inherent tumor complexity and molecular heterogeneity, it is well evidenced that patients with similar pathological and diagnostic cancer phenotypes often respond differently to the same treatment [1, 2]. Thus, this has spurned efforts to transition from traditional one-size-fits-all approach to precision or personalized medicine – allowing clinicians to match the right drug to the right patient [3]. Precision medicine and targeted therapies are progressing rapidly as scientists develop a deeper understanding of patients’ genome-level heterogeneity as well as prognostic and diagnostic biomarkers [4–8]. There are two related axes of investigation that have contributed to these efforts: pre-clinical model systems and high-throughput molecular profiling techniques; as we expound below.

Cell lines are samples taken from human tumors and cultivated in labs, serve as fundamental preclinical models for studying the pathological mechanisms and treatment efficacy of cancer [9]. They are pivotal because they offer a controlled approach to discover underlying biological mechanisms and hence, effective cancer treatments [9]. However, accumulating evidence suggests that inherent biological differences between preclinical models and patient tumors systematically hinder translational research efforts – from bench-side knowledge to its use at the clinical bedside [10–13]. Specifically, from a genomics perspective, cell lines can genetically drift over prolonged culture periods, potentially adopting mutations and traits that render them less representative of the original patient tumors [11]. This challenge underscores the need for a systematic and comprehensive examination of the similarities and differences between preclinical and clinical (i.e. patient tumors) model systems to uncover novel drug targets and prediction models in cancer.

The advances in large-scale genomic and pharmacological profiling across patient tumors and preclinical models make this investigation possible. The sources of patient tumor datasets such as The Cancer Genome Atlas (TCGA) [14], the International Cancer Genome Consortium (ICGC) [15] and The Cancer Proteome Atlas (TCPA) [16] have provided extensive multi-platform genomics (multi-omics) profiles of thousands of cancer and normal samples spanning > 30 cancer types. The data resources of preclinical model systems, including Cancer Cell Line Encyclopedia (CCLE) [17], and MD Anderson Cell Lines Project (MCLP) [18] contain multi-omics data of > 1, 000 cell lines. The large-scale drug perturbation studies of biological model systems such as Genomics of Drug Sensitivity in Cancer (GDSC) [19], Dependency Map (DEPMAP) [20], Library of Integrated Cellular Signatures (LINCS) [21], and Connectivity Map (CMAP) [21] have characterized their responses to a wide range of anti-cancer therapeutics. These datasets enable the assesment of the translational potential of cell lines to patient tumors, by identifying their differential and shared genomic characterization and its effect in drug response prediction.

To this end, computational and clinical researchers have made significant progress in therapeutic biomarker detection and drug response prediction by using cell lines alone. In the context of biomarker detection, high-throughput genomic studies have examined the correlation between drug response with mutations and copy number alterations based on cell lines [22]. With respect to drug response prediction, statistical models have been trained solely on cell lines’ genomic and drug perturbation data, and tested on out-of-sample data from cell lines or patient tumors [23–25]. The data from cell lines have been widely used for model training because high-throughput drug response data across hundreds of cell lines for many drugs are publicly available in large databases, such as the aforementioned CCLE and GDSC databases. On the other hand, since clinical data of patients are often very expensive and time-consuming to generate, these data are still relatively small in sample size and present challenges to be used as reliable training data for drug response prediction [26].

Even though cell lines have been typically used to train drug response prediction models, the *in vitro* response from cell lines is not expected to adequately represent *in vivo* response from primary patient tumors [27, 28]. The main reason is that cell lines evolve quite differently from patient tumors in the absence of the complex *in vivo* microenvironment, and genotypic and phenotypic variation in cell lines can happen over a large number of passages [29, 30]. Thus, given *in vivo* drug response prediction as the primary goal, the variation between cell lines and patient tumors makes the models, which are trained solely on cell lines, suffer from the fundamental problem that training and test sets come from two different biological model systems [31].

To mitigate the effect of the differences between biological model systems on drug response prediction, genomic data from patients have been used in model training to integrate genomic features between cell lines and patients. Previous drug response prediction methods [32–34] assumed that batch effects were the main source of differences between cell lines and patients’ gene expression data. More recently, some methods [31, 35–37] have employed domain adaptation technique, a subcategory of transfer learning, to bridge the gap between preclincial models and patient tumors by identifying a shared latent space common to both the cell lines’ and patients’ domains. However, these methods do not explicitly assess the model-specific structured variation in gene expression data (pertaining to either cell lines or patients) during model construction. In such instances, if the modeling of gene expression data for cell lines and patients solely comprises shared latent structure and residual noise, the model-specific information will be incorporated into the estimated shared latent structure and/or residual matrix. We hypothesize (and show) that, this could influence the estimation accuracy of the shared structure, and subsequently affect the prediction of drug responses using the estimated shared variation.

To this end, we propose an integrative principal component regression (iPCR) model that serves four key purposes: 1) to quantify the similarities and differences in gene expression data between cell lines and patient tumors, 2) to use the patient-cell line similarities to predict the patient drug response, 3) to find key genomic drivers (e.g., pathways, gene sets) for patient drug-specific response, and 4) to infer model-specific and shared co-expression networks between cell lines and patients and detect hub genes in them. Viewed as a domain adaptation model, our innovative approach not only adapts multi-view statistical techniques to address the unique challenges of multi-system prediction problem in precision oncology, but also comprehensively investigates the common biology and differences between cell lines and patient tumors. This approach provides enhanced interpretability of both gene effects on drug response and gene-gene interactions, offering a fresh perspective in this critical domain. Figure 1 shows the overall conceptual flow of our pipeline. In stage I of iPCR model (Figure 1A), we integrate gene expression data between cell lines and patients, and identify latent shared features (i.e., joint principal scores) and model-specific features (i.e., individual principal scores). These latent features essentially capture the underlying shared and model-specific biological process across the two model systems. In stage II of iPCR model (Figure 1B-C), the shared features from stage I and drug responses of cell lines are used to train a penalized regression model for the prediction of patient drug responses. Finally, the joint and individual variations across the two biological model systems detected by the iPCR model are used to capture the model-specific and shared gene co-expression networks between the biological model systems.

**Fig. 1.**
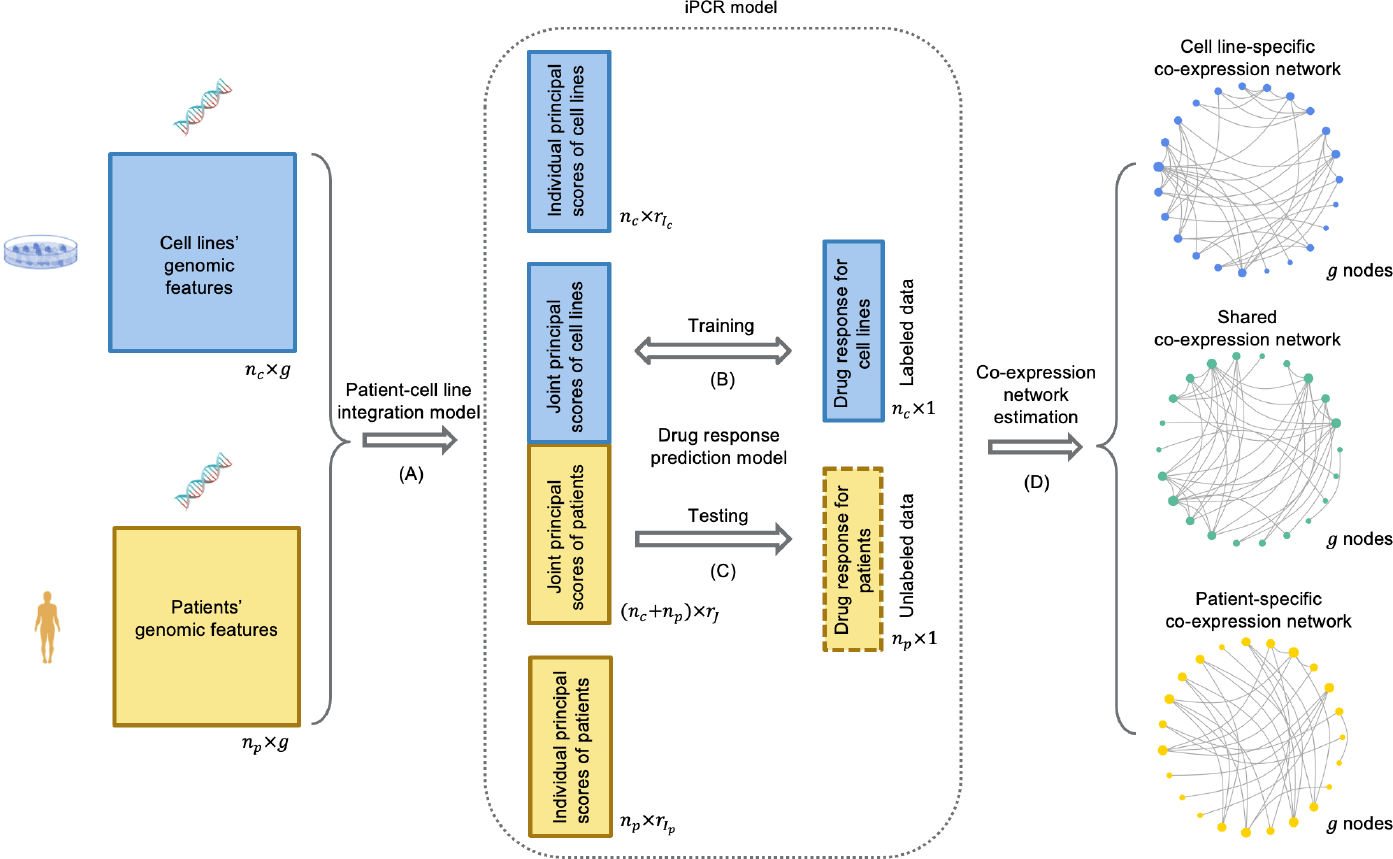
Conceptual framework of the iPCR model. The joint model is specified by two sub-models, namely the patient-cell line integration model (A) specifying the biological similarity (joint principal scores) and difference (individual principal scores) between cell lines and patients, and the drug response prediction model (B and C) specifying the relationship between drug response and the joint principal scores of cell lines and patients, which is trained on the labeled cell lines’ data and tested on the unlabeled patients’ data. The results of iPCR model are then used to infer model-specific and shared co-expression networks through co-expression network estimation (D).

We demonstrate that iPCR outperforms alternative approaches in patient drug response prediction, under multiple simulated scenarios and multiple cancer drugs in a pan-cancer application. By implementing our iPCR and gene co-expression network decomposition methods, we identify key genomic drivers (e.g., pathways, gene sets) of patient drug-specific responses, and hub genes in the shared co-expression network between cell lines and patients.

The rest of this manuscript is organized as follows: Section 2.1 introduces the formats of genomic and drug response data used in our study. Section 2.2.1 formulates the stage I of iPCR, which employs a dimension reduction technique to decompose multi-system genomic data of cell lines and patients into shared and individual components. Section 2.2.2 describes the stage II of iPCR, which utilizes the outputs from stage I to predict the drug response in patients based on cell lines’ drug response data. Section 2.3 delineates the details of iPCR model estimation. Section 2.4 introduces the gene co-expression network decomposition method based on the outputs from the iPCR model. Section 3 evaluates the prediction performance of iPCR and its sensitivity to rank selection through simulation studies. In Section 4, we apply iPCR and gene co-expression network decomposition methods on large-scale pan-cancer data from cell lines and patient tumors. We conclude with a discussion in Section 5.

## 2 Methods

### 2.1 Data formats

Following the conceptual framework in Figure 1, the specific data formats are defined as follows. We consider two gene expression matrices 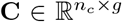 from *n*_*c*_ cell lines with *g* gene features, and **P** ∈ 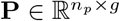from *n*_*p*_ patients with the same set of *g* gene features. Then, these two matrices can be combined into a single data matrix 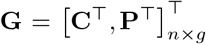, where *n* = *n*_*c*_ + *n*_*p*_ is the total number of samples. Following standard preprocessing in [37, 38], we performed a centralization and standardization for each column of **C** and **P**.

For a given drug *D* assayed on *n*_*c*_ cell lines, we denote vector 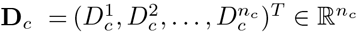 as the observed continuous drug responses for those cell lines. For the same drug taken by *N*_*p*_ patients, the vector 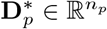 contains the drug responses for those patients, where (*) represents the unobserved labels. Our primary objective is to predict unobserved patient drug responses 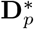 based on the available datasets {**C, P, D**_c_ } . In this paper, **D**_*c*_ is measured by the area under the drug response curve (drug-AUC), although other omic platforms (e.g., proteomics, metabolomics) and drug response metrics (e.g., IC50) can easily be used in our framework.

### 2.2 iPCR model

In this section, we detail the two stages of our iPCR model. We first formulate the patient-cell line data integration model based on variation decomposition, which helps identify shared patterns of variation (called joint structure) and model-specific variation (called individual structure) between the gene expression data of cell lines and patients. Then, we introduce the drug response prediction model, which takes advantage of the latent shared features between patients and cell lines from the joint structure, to predict the drug responses in patients and identify the key therapeutic biomarkers.

#### 2.2.1 Stage I: patient-cell line integration model

Our proposed patient-cell line data integration model assumes the existence of both (genomically driven) biological similarity and difference between the two model systems. Based on this assumption, the model decomposes **C** and **P** into a low-rank joint structure that captures the common features between cell lines and patients; two low-rank individual structures that preserve individual features that are not shared, and the residual noise. The model is formalized as

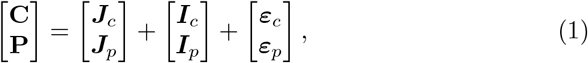

where for all *i ∈ {c, p}*, ***J***_*i*_ is the sub-matrix of the joint structure matrix 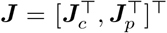 with rank *r*_*J*_, ***I***_*i*_ is the individual structure matrix with rank 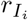, and ***ε***_***i***_ is the error matrix containing the independent entries with 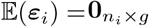. We denote *r*_*J*_ as joint rank, 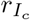as individual rank of cell lines, and 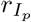 as individual rank of patients.

Model 1 necessitates the conditions detailed in Proposition 1 for unique identifiability, as follows:

##### Proposition 1

*[38, 39] Let* row(*·*) *and* col(*·*) *be the column and row space of a matrix, respectively. Given a set of matrices {****G***_*i*_, *i=c, p}, there are unique sets of matrices {****J***_*i*_, *i = c, p} and {****I***_*i*_, *i = c, p} such that*

1. ***G***_*i*_ = ***J***_*i*_ + ***I***_*i*_ *for i = c, p*
2. col(***J***_*i*_) *∩* col(***I***_*i*_) = 0 *for i = c, p*
3. row(***J***_*i*_) = row(***J***) *for i = c, p, where* 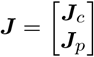
4. row(***J***_*i*_) ⋂ row(***I***_*i*_) *for i = c, p*
5. row(***I***_*c*_) *∩* row(***I***_*p*_) = 0

Intuitively, these conditions suggest that gene expression data from either cell lines or patients can be decomposed into a joint structure and an individual structure, excluding noise (Condition 1). Moreover, the data from cell lines and patients share a latent common feature space within the joint structure (Condition 3), and this shared feature space is linearly independent of the individual feature space of either patients or cell lines (Condition 4). Additionally, the individual structure of patients doesn’t share a feature space with the individual structure of cell lines (Condition 5). This aligns well with our assumption about the existing similarities and differences in gene expression data between cell lines and patients. As mentioned in [39], Condition 2 on the column spaces is equivalent to the rank condition in [38], which can be expressed as *rank*(***J***_*i*_ + ***I***_*i*_) = *rank*(***J***_*i*_) + *rank*(***I***_*i*_) for *i* = *c, p*. This condition is necessary for the uniqueness of the decomposition at Stage I, in conjunction with Condition 4.

To fit the various components of Model 1, we employ PCA or singular value decomposition (SVD) on the joint structure ***J***, as well as the two individual structures ***I***_*c*_ and ***I***_*p*_. To this end, Model 1 can be further factorized as follows:

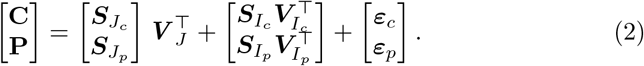

The joint score matrix 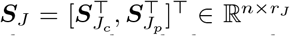 contains *r*_*J*_ joint principal scores from ***J***, which can be viewed as the latent shared features of cell lines and patients, capturing their biological similarity. The orthonormal joint loading matrix 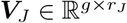quantifies the extent to which each original feature in ***J*** is related to the latent shared features, such that 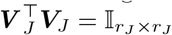. From another angle, the columns of ***V***_*J*_ (i.e., shared principal axes) define an *r*_*J*_ –dimensional shared feature space between cell lines and patients, while each row of ***S***_*J*_ provides the location of the corresponding sample in this shared feature space. The score matrices 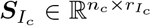 and 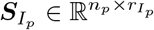 summarize the features individual to cell lines and patients, with the corresponding loadings 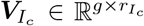 and 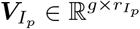, respectively.

##### Relation to JIVE

The model formalization above is similar in some respect to the JIVE model proposed by Lock et al [38], since both decompose the original unaligned data into joint and individual structures as well as an error matrix. However, the difference is that JIVE and its related methods [39–41] are designed to integrate multi-source data (i.e., the same set of samples with data from different sources), while iPCR aims to integrate multi-system data (i.e., different biological model systems with the same set of features). Consequently, in the JIVE model, the two input matrices share the same sample count, but in our model, **C** and **P** share the same gene count. This difference implies that the joint loading matrix 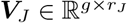 in our Model 2 corresponds to a joint score matrix 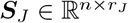in JIVE, leading to different interpretations. Here, *n* represents the common sample size in JIVE’s multi-source context.

#### 2.2.2 Stage II: Drug response prediction model

At this stage, we use the estimated 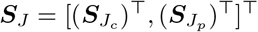 from Stage I, which captures the shared biology between cell lines and patients, along with the observed **D**_*c*_, to predict the unobserved 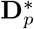 using a penalized linear regression model. We choose a penalized model ranther than a non-penalized one because our pan-cancer application in Section 4 is in a high-dimensional setting, and the value of *r*_*J*_ may not be very small regarding the sample size. For example, in our real application, we choose *r*_*J*_ = 55 for most drugs through rank selection. To avoid overfitting, in this paper, we use the elastic net regularization for the estimation of Model 3. We note here that the penalized Gaussian linear regression model suffices for our case study and is easy for interpretation, although any other linear or nonlinear prediction models could be potentially used in this stage.

Specifically, the drug responses are regressed on the *r*_*J*_ shared features in ***S***_*J*_ from Stage I, rather than the original *g* gene features. The drug response model for cell lines can be formularized as follows:

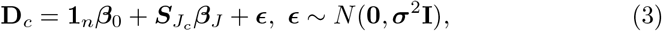

where **1**_*n*_ is a *n ×*1 vector of ones, ***β***_0_ is a constant intercept, ***β***_*J*_ is a *r*_*J*_*×* 1 vector of regression coefficients that measure the effect of the corresponding shared features on drug responses, and ***ϵ*** is a *n ×* 1 error vector. Since 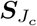 and 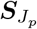 capture the shared biology between the two biological model systems, we assume that the same latent shared feature in both 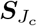 and 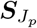 has an identical effect on their drug responses. Based on this assumption, we can then use 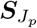, the estimated ***β***_0_ and the estimated ***β***_*J*_ to predict the unobserved patient drug responses 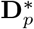 as detailed in Section 2.3.1.

To enhance the interpretability of regression coefficients, it is essential to transition from the latent feature space to the gene space, which offers clearer biological significance. Specifically, 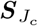 can be approximated by projecting the original cell lines’ gene expression data onto the latent shared feature space such that 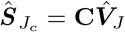 . By substituting this into Equation 3, we can get

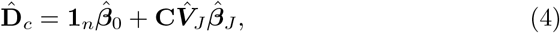

where 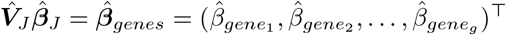 represents *g×* 1 vector of estimated regression coefficients for shared gene effects between cell lines and patients on drug responses. In addition, the ranked list of these gene effects can be used as input in the PreRanked gene set enrichment analysis [42].

### 2.3 Model estimation

In this section, we outline the estimation process for our two-stage iPCR model. First, we discuss the estimation of latent joint and individual structures at Stage I, given their predetermined ranks. Next, we explain how the model training and drug response prediction are conducted at Stage II, using shared features between cell lines and patients derived from the joint structure. Then, we describe the method for rank selection, optimized through cross-validation. Lastly, when the sample sizes of cell lines and patients differ significantly, we introduce a scaling technique at Stage I to balance the data integration process, followed by a corresponding re-scaling at Stage II.

#### 2.3.1 Stages I and II given joint and individual ranks

For the estimation of patient-cell line integration model at Stage I (Model 1), suppose the joint rank *r*_*J*_ and individual ranks 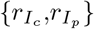 are given. As mentioned in Section 2.2.1, although our patient-cell line integration model and JIVE have different problem settings and interpretations, their model formulations are similar. Hence, we use the algorithm introduced by [38] and “r.jive” package [43] to estimate the joint and individual structures with the given ranks 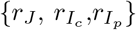. The final estimated joint and individual structures are denoted as ***Ĵ, Î***_*c*_, and ***Î***_*p*_. Then, SVD is performed on ***Ĵ*** such that 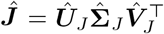. Those *r*_*J*_ estimated shared features between cell lines and patients can be acquired from the estimated joint score matrix 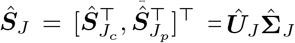, which allows us to use Model 3 to predict patient drug response given the responses from cell lines to a certain drug *D*.

Given ***Ŝ***_*J*_ from Stage I, we use **D**_*c*_ and 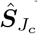as the training data to fit Model 3 with an elastic net regularization [44]. The regression coefficients 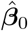 and 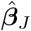 are shrunk based on both L1-norm and the L2-norm. The elastic net model training is done in R based on the caret package [45]. Finally, the predicted responses of *n*_*p*_ patients to the given drug *D* are given by

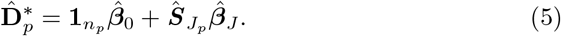

#### 2.3.2 Rank selection of joint and individual structures

The unsupervised rank selection methods [38] tend to neglect those latent signals that explain a very small proportion of variance in the data, although they could be strong predictors of outcomes. This could result in underestimation of the true rank and subsequently impact prediction accuracy. Hence, analysts sometimes select the optimal number of principal components based on the prediction performance through cross-validation in the labelled source data [46, 47]. In addition, a domain adaptation study [48] showed the advantage of using label information in the source domain data to improve prediction performance. Inspired by these studies, we select an optimal rank set 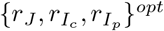 by performing a 10-fold cross-validation based on **D**_*c*_ and 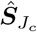for a certain drug.

To provide more details for Algorithm 1, we first choose *m* candidates of rank set {**r**_1_, …, **r**_*m*_} for grid search, such that 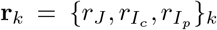, where *k* = 1, …, *m*. Since the number of combinations of 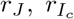 and 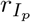 grows cubically with the number of candidates for each type of rank, the exhaustive grid research is only feasible for low-dimensional training data. Due to the high dimension of pan-cancer genomic data, we limit the number of combinations by only doing a grid search on *r*_*J*_ . Given a certain *r*_*J*_, the corresponding 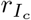 and 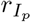are calculated by subtracting *r*_*J*_ from the estimated ranks of denoised **C** and **P**, respectively; the rank of denoised **C** or **P** is estimated based on the inflection point of the cumulative eigenvalues through PCA, or, if there is no obvious inflection point, the parallel analysis [49] through “paran” package [50].

##### Algorithm 1 iPCR model

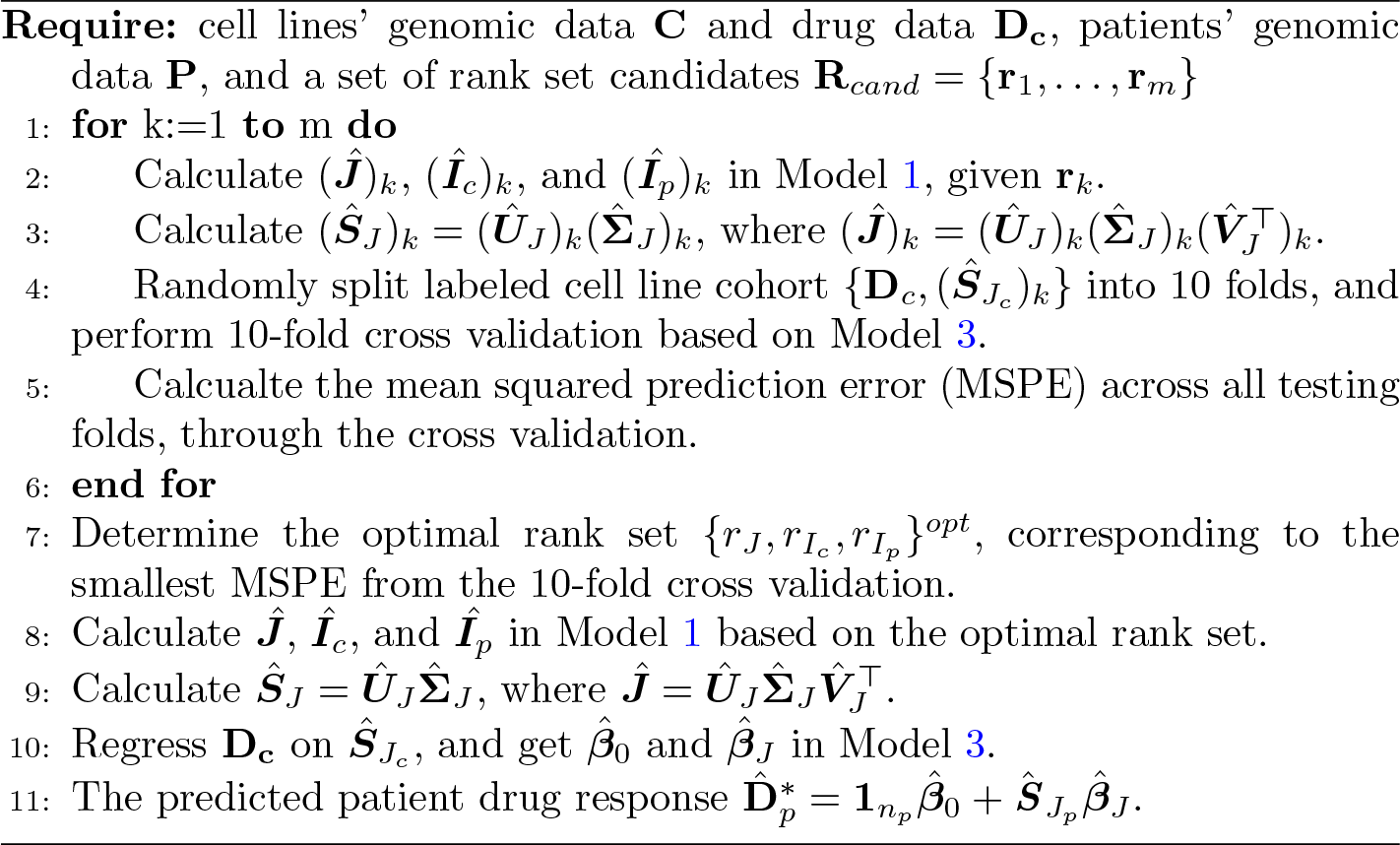

As Algorithm 1 shows, for each candidate rank set **r**_*i*_, we then 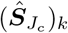 from the patient-cell line integration model, and randomly split labeled cell line cohorts 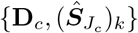 into 10 folds. Next, we perform 10-fold cross validation based on Model 3, and calculate the mean squared prediction error (MSPE) across all testing folds corresponding to **r**_*k*_. Last, the optimal rank set 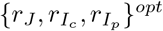 is chosen with the smallest MSPE from the cross validation. The optimal rank set is used for the training of the iPCR model to predict 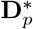.

#### 2.3.3 Data scaling at Stages I and II

In scenarios where the sample sizes in **C** and **P** exhibit substantial discrepancies, “the bigger dataset always wins” during data integration at Stage I. This bias can adversely affect the accuracy of predictions at Stage II. To avoid this issue, it is helpful to scale **C** and **P** by their total variation such that 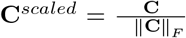, and 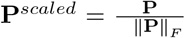, so that 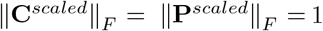, where ∥·∥_*F*_ defines the Frobenius norm [38, 39]. At Stage I, using the scaled matrices **C**^*scaled*^ and **P**^*scaled*^ as inputs, we obtain the estimated scaled joint score matrices 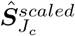 and 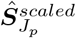 following the methodology in Section 2.3.1. At Stage II, these matrices are scaled back such that 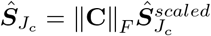 and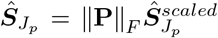, which are subsequently utilized for model training and patient drug response prediction.

### 2.4 Gene co-expression network estimation

Genes are naturally interconnected through biological networks [51]. Various network-based models have been developed to identify gene co-expression networks, aiming to yield insights into underlying biological and regulatory pathways [51–55]. However, it is not feasible to assume a universal network structure across different types of biological samples [56]. Acharyya et al. [56] proposed a model, known as SpaceX, to discern both shared and cluster-specific gene co-expression networks by decomposing the covariance matrix of genes into its shared and cluster-specific components. Inspired by this approach, in this section we provide a new angle to study the gene co-expression network in cell lines and patients, based on Model 1, which is also a byproduct of iPCR. Unlike SpaceX, which relies on a factor model and decomposes the true underlying covariance matrix, iPCR is related to PCA and focuses on decomposing the sample covariance matrix.

The estimated denoised gene expression matrices for cell lines and patients 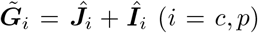 from step 8 in Algorithm 1 are used as input herein this section. Let |·| be the cardinal number of a set. We define the gene coexpression network as a graph {*V, E}*, where the vertex set *V* denotes *g* genes with *V* = *g* and *E* represents the set of undirected pairwise relationships (edges) between the *g* genes such that |*E*| = *g*(*g −* 1)*/*2. The edge between two gene vertices denotes their co-expression level, which is defined using a similarity measure, Pearson correlation coefficient. Our gene co-expression network estimation method decomposes the gene network into two hierarchical components: 1) a shared component representing the shared gene co-expression network across the two biological model systems (cell line and patient); and 2) a model system specific component representing the gene co-expression network individual to the cell lines or patients.

For all *i* ∈ {*c, p*}, we propose a sample covariance decomposition method for 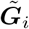, such that

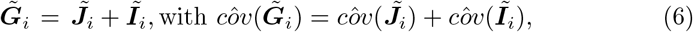

where 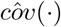 represents sample covariance matrix between features (i.e., genes). The matrices 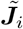and 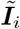can be uniquely determined under the conditions specified in Proposition 2. The unique solution and the proof of identifiability are provided in Supplementary Material Section S1.1 and Section S1.2, respectively. Note that Propositions 1 and 2 have the same Condition 3, that there is a shared feature space between cell lines and patients. The only difference between them is that in Proposition 1 the row spaces between ***J***_*i*_ and ***I***_*i*_ are orthogonal, while Proposition 2 has the constraint that column spaces between 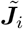and 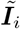 are orthogonal, for all *i* ∈ {*c, p*}. Condition 2 of Proposition 2, which focuses the orthogonality of column spaces, serves as a key constraint that ensures 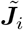 and 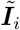satisfy the property of sample covariance decomposition in Model 6. Further, Condition 4 in Proposition 2 is essential for the uniqueness of the decomposition. When paired with Condition 2, it also guarantees the property such that 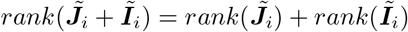.

#### Proposition 2

*Given a set of matrices* 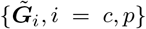, *there are unique sets of matrices* 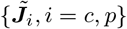 *and* 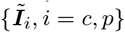 *such that*,

1. 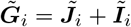*for i = c, p*
2. 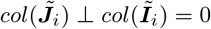 *for i = c, p*
3. 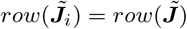 *for i = c, p, where* 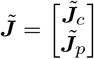
4. 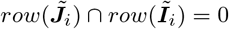 *for i = c, p*
5. 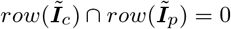

For easier interpretation, for all *i* ∈ {*c, p*}, the sample covariance matrices 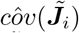 and 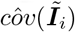 are transformed to the sample correlation matrices 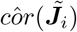 and 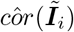, respectively. For example, the (*m, n*)^*th*^ element of 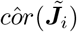, denoted as 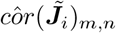, represents the shared sample correlation between *m*^*th*^ gene and *n*^*th*^ gene in the cell lines’ gene expression data (*i* = *c*) or patients’ gene expression data (*i* = *p*). An edge between two genes is considered to be significant in the co-expression network if the corresponding correlation’s absolute value is larger than the threshold *δ* (e.g., *δ* = 0.7). To be conservative, we define the edge between *m*^*th*^ gene and *n*^*th*^ gene as significant in the shared gene co-expression network between cell lines and patients, if both 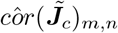 and 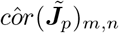 are larger than *δ*. We then use the shared and model-specific gene co-expression networks to detect hub genes based on the number of edges linked to each gene.

## 3 Simulation Study

We compare the drug response prediction accuracy of our iPCR model with other alternative methods in the simulated cell line and patient data mimicking our real data applications (Section 4). The simulated data are generated under different levels of similarity between cell lines and patients and signal-to-noise ratios. We also evaluate the sensitivity of rank selection relative to the prediction performance for iPCR.

### 3.1 Simulation design and comparative analyses

This simulation study examines the performance of iPCR and other methods under different levels of similarity between cell lines and patients and degrees of noise. We assume the dimensions of **C** and **P** are 1500*×* 2000 and 10000 *×*2000, respectively, where rows correspond to samples and columns correspond to genes. **C** and **P** are sampled based on the following models:

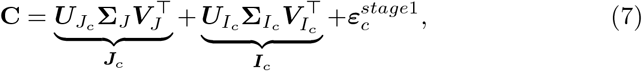

and

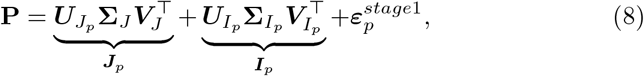

where 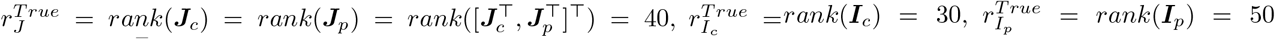, the elements of 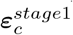follow 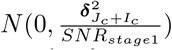 and the elements of 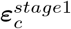follow 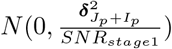. For all *i ∈ {c, p}*, the signal-to-noise ratio *SNR*_*stage*1_ is defined as the ratio between the sample variance of elements in the denoised data ***J***_*i*_ +***I***_*i*_ (denoted as 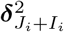) and the variance of 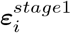. The shared feature space *row*(***V***_*J*_) is orthogonal with the individual feature spaces 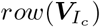 and 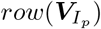, and 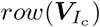 is also orthogonal with 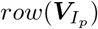. We use the term “joint proportion” to refer to the ratio of 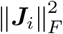 to 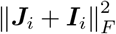, which quantifies the proportion of variation of ***J***_*i*_ relative to the denoised genomic data ***J***_*i*_ + ***I***_*i*_ for all *i* ∈ {*c, p*}. In other words, a larger value of joint proportion corresponds to a higher level of similarity between cell lines and patients. In order to meet orthogonality constraints and control different levels of joint proportion from 20% to 80%, the SVD components of ***J***_*c*_, ***J***_*p*_, ***I***_*c*_, and ***I***_*p*_ in Model 7 and 8 are constructed separately (more details in Supplementary Material Section S2.1). We also set different levels of *SNR*_*stage*1_ at 0.5, 1, or 2.

We assume that **D**_*c*_ and **D**_*p*_ depend on the joint score matrices 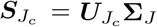and 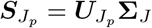, respectively, such that

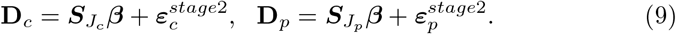

The regression coefficient ***β*** is a vector of length 40, where the elements are generated from *N* (0, 4). The elements of 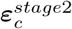 and 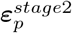 follow 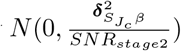 and 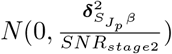, where 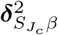 and 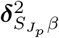 represent the sample variances of 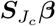 and 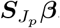, respectively. The signal-to-noise ratio in Model 9, *SNR*_*stage*2_, is set to 5. The prediction performance is measured by the Spearman correlation between the true and predicted values of **D**_**p**_, focusing solely on the order of values. In each setting, we generate 50 replicates of {**C, P, D**_c_} for training and **D**_*p*_ for testing.

For the iPCR model, six candidate rank sets are chosen for grid search. Specifically, the candidate rank sets 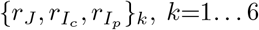, include {10, 60, 80}_1_, {20, 50, 70}_2_, {30, 40, 60}_3_, {40, 30, 50}_4_, {50, 20, 40}_5_, and {60, 10, 30}_6_ based on the rank selection guideline in Section 2.3.3. We compare with 1) an elastic net regression trained solely on cell line data (Elastic Net); 2) an elastic net regression trained and tested on the batch-corrected data (Combat + Elastic Net), similar to [33]; and 3) an elastic net regression trained and tested on the first *K* principal scores of [**C**^*⊤*^, **P**^*⊤*^]^*⊤*^ (*K* selected through 10-fold cross validation), which in this paper is called principal component regression PCR). Both PCR and iPCR aim to detect the joint structure based on variation decomposition, but PCR does not consider individual structures during the modeling. To make an apples-to-apples comparison, we consistently use Elastic Net as the regularization method across the above methods.

Figure 2 demonstrates the prediction accuracy (measured by the Spearman correlation between the predicted and true **D**_**p**_) of different methods over 50 replications for each combination of *SNR*_*stage*1_ and joint proportion. For better illustration, we plot the curves of Spearman correlation as a function of joint proportion in three subplots that correspond to three levels of *SNR*_*stage*1_. As expected, when joint proportion increases, the prediction accuracy of all methods increases, since the expected drug responses are assumed to only depend on the similarities between cell lines and patients. In addition, a larger *SNR*_*stage*1_ leads to a higher prediction accuracy for all methods. iPCR outperforms the competing methods under different levels of *SNR*_*stage*1_ and joint proportion. Surprisingly, Combat + Elastic Net performs very similarly with Elastic Net, which means that the Combat batch removal method does not filter model-specific differences in our setting. It is also fair to say that the batch effect is negligible in this instance. PCR performs worse than iPCR, in part because in PCR the first *K* principal scores used for prediction includes both shared and model-specific information. As joint proportion increases, those first *K* principal scores in PCR include more information from the joint structure, so it performs closer to iPCR in terms of the prediction accuracy. When *SNR*_*stage*1_ = 0.5, PCR always outperforms Elastic Net and Combat + Elastic Net at different levels of joint proportion. For *SNR*_*stage*1_ = 1 (or 2), PCR only outperforms Elastic Net and Combat + Elastic Net when joint proportion is larger than 0.6 (or 0.7); possibly because Elastic Net utilizes all the data including the information of similarity in prediction while PCR has difficulty detecting the information of similarity when joint proportion is low.

**Fig. 2.**
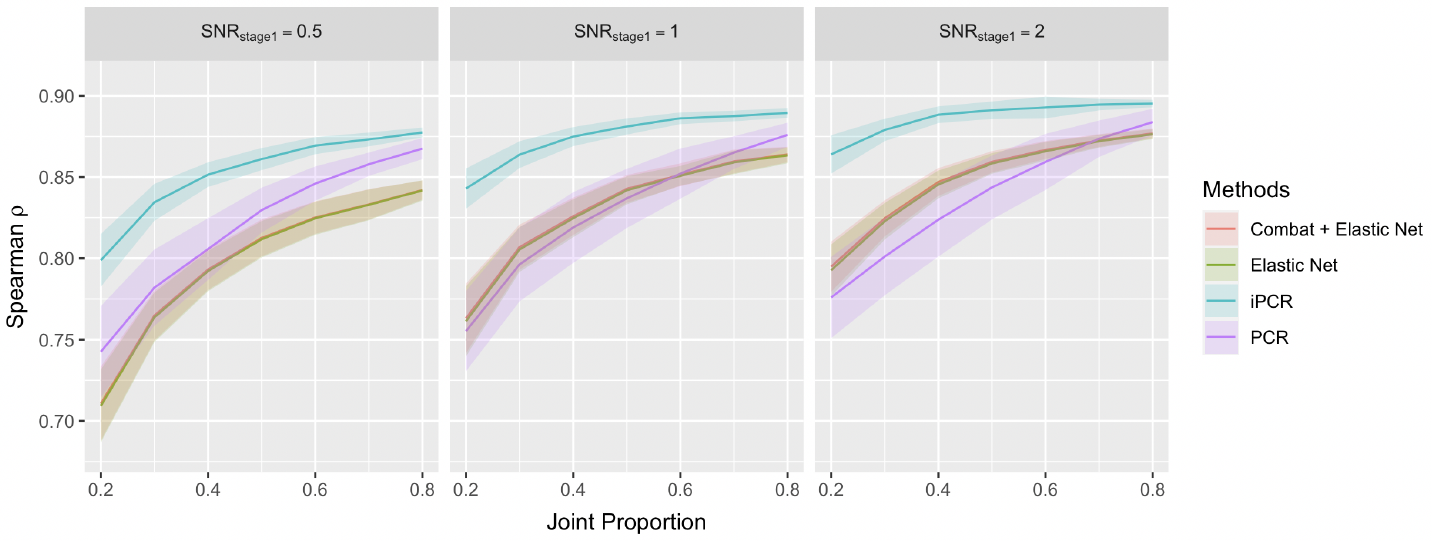
Accuracy comparison of different methods in predicting **D**_**p**_ in simulation study. The prediction performance is measured by Spearman correlation between the true and predicted values of **D**_**p**_. The data are generated under different levels of *SNR*_*stage*1_ and joint proportion over 50 replications. The sample mean of Spearman correlation over replications correspond to the solid curves, and the shaded areas represent mean *±* 1sd.

### 3.2 Sensitivity analysis of rank selection for iPCR

To measure the sensitivity of rank selection to the prediction accuracy for iPCR, we implement iPCR given a certain rank set without doing the rank selection, following the data generating process in Section 3.1. The rank sets of interest in the form of 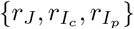 include 1) {40, 30, 50} that corresponds to the true rank set 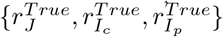; 2) {30, 40, 60} and {50, 20, 40} that under- and over-estimate 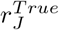, respectively, while the denoised ranks of **C** and **P** are correctly chosen; and 3) {40, 20, 40} and {40, 40, 60} that under and over estimate the true individual ranks, respectively, while the joint rank is correctly chosen. The accuracy comparison of iPCR given different rank sets in predicting **D**_**p**_ is shown in Figure S1. We note that the accuracy curve of iPCR in Figure 2 is very similar to the accuracy curve of iPCR given the true rank set in Figure S1. This observation illustrates that in Section 3.1 iPCR always correctly selects the true rank set for prediction; although it sometimes does not select the true rank set, it achieves comparable accuracy to iPCR given the true rank set. In addition, given the four misspecified rank sets defined in this section, iPCR still outperforms the competing methods in Section 3.1, except iPCR given {30, 40, 60} (the case that under estimates 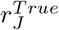) when joint proportion is high, as shown by Figure 2 and S1. Further discussion of the sensitivity analysis of iPCR rank selection can be found in Supplementary Material Section S2.2.

## 4 Application to pan-cancer drug response data

We performed a retrospective in-silico analysis to evaluate our iPCR model on a pan-cancer dataset. Our primary scientific goals are to: 1) identify top drugs based on prediction performance on patient tumors using cell-line data; 2) infer and interpret iPCR gene signatures associated with drug responses to identify novel drug targets; and 3) investigate the joint and individual gene coexpression networks between cell lines and patients, to identify co-ordinated mechanisms underlying drug resistance.

### 4.1 Dataset description

All datasets used in our application are publicly available. Gene expression data for 12,747 patient tumors from >40 cancer types were downloaded from the Tumor Compendium v11 Public PolyA, a database maintained by The Treehouse Childhood Cancer Initiative at the UCSC Genomics Institute [57]. These patient tumors include samples from Treehouse clinical sites, the Therapeutically Applicable Research to Generate Effective Treatments (TARGET) project, and The Cancer Genome Atlas (TCGA) [14]. Gene expression data for 1,406 cell lines from 30 different tissues were obtained from the DepMap Public 22Q2 file, named as “CCLE expression full.csv” [17]. The gene expression data for both patient tumors and cell lines were pre-processed using established pipelines, as described in [17, 57]. Additionally, the data scaling pipeline outlined in Section 2.3.3 was applied due to the large difference in sample size between patients and cell lines. We restricted our analysis to 1,774 cancerrelated genes [58] that were present in both tumor and cell line data, and performed a gene-level centralization and standardization.

The *in vivo* drug response data of cell lines were obtained from the Genomics of Drug Sensitivity in Cancer (GDSC) panel [19]. The *in vitro* response data of patients to 13 different cancer drugs for validation instead of model fitting, were obtained from [59]. The drug response data of cell lines and patients contained a subset of samples in the gene expression data. The number of patients with available drug response data for each drug is listed in Table 1, while the cell line sample size exceeds 600 for each drug. For cell lines in the GDSC panel, drug response was measured by the area under the drug response curve. This metric integrates information about both drug efficacy and potency into a single value, and has been previously reported as the most robust metric [60]. Note, to distinguish the name of the drug response metric with the area under the ROC curve (AUC) used for measuring prediction performance, we named the former as the drug-AUC. Large values of drug-AUC are associated with drug resistance. We classified patient drug responses into two categories: Response (including “Complete Response” and “Partial Response”) and Non-response (including “Stable Disease” and “Progressive Disease”).

**Table 1.** Performance results, as measured by AUC, for iPCR and four competing methods in patient drug response prediction.

| Drug | Patient Samples | Elastic Net | Combat + Elastic Net | PCR | TRANSACT | iPCR |
| --- | --- | --- | --- | --- | --- | --- |
| Trastuzumab (Afatinib) | 16 | 0.82 | 0.82 | **1.00** | 0.71 | **1.00** |
| Etoposide | 86 | 0.52 | 0.53 | 0.41 | 0.68 | **0.70** |
| Vinorelbine | 31 | 0.60 | 0.61 | 0.64 | 0.64 | **0.67** |
| Cisplatin | 309 | 0.68 | **0.68** | 0.67 | 0.64 | 0.66 |
| Oxaliplatin | 38 | 0.57 | 0.58 | 0.59 | 0.63 | **0.65** |
| Gemcitabine | 158 | 0.64 | 0.64 | 0.63 | **0.65** | 0.61 |
| Temozolomide | 94 | 0.51 | 0.50 | **0.68** | 0.50 | 0.58 |
| Pemetrexed | 39 | 0.57 | 0.57 | **0.65** | 0.42 | 0.56 |
| Paclitaxel | 157 | 0.50 | 0.51 | 0.54 | **0.60** | 0.53 |

*Note:* The drug name in the parenthesis corresponds to the matching drug from GDSC. For each drug and method, we compared the predicted continuous patient drug response to the known binary clinical response. We reported patient sample size for testing and AUC for measuring the prediction performance. Bold cells correspond to the largest AUC across the five methods for predicting the patient response to a certain drug.

### 4.2 Patient drug response prediction

To validate the prediction performance of our iPCR model, we first aligned the whole cell line and patient expression data based on stage I of iPCR (Section 2.2.1). Next, for each drug, we used the cross-validation method described in Section 2.3.3 to select the best combination of joint and individual ranks among the candidates. Following the rank selection guideline in Section 2.3.3, we chose 20, 25, …, 60 as the candidates for joint ranks. For each candidate joint rank, the corresponding individual ranks of cell lines and patients were obtained by subtracting the joint rank from 64 and 84 (the estimated ranks of denoised cell line and patient gene expression data based on parallel analysis [50]), respectively. Then, for each drug, given the optimal ranks, we trained the regression model in stage II of iPCR (Section 2.2.2) to predict the drug-AUC values for patients. Finally, we calculated the AUC by comparing the predicted drug-AUC values between known responders and non-responders. The higher the AUC, the better the prediction performance.

The competing approaches for patient drug response prediction included the three alternative methods mentioned in Section 3.1, and TRANSACT [37]. TRANSACT is a state-of-art domain adaptation approach for drug response prediction, which trains an elastic net regression model based on the aligned kernel principal components of cell line data. As recommended by [37], we used *γ* = 5 *×* 10^*−*4^ (amount of non-linearity), which is a tuning parameter of TRANSACT calibrated through the transfer learning from cell lines to PDX models. Note that during model training, except for Elastic Net regression, all other methods utilized the patient gene expression data, while none of the methods used patient drug responses.

In Table 1, we show the prediction performance for 9 drugs that exist at least one approach with AUC > 0.6. Cells in bold indicate the highest AUC among the five methods for predicting the patient response to a certain drug. Our iPCR approach outperforms other methods in 4 drugs (Trastuzumab, Etoposide, Vinorelbine, and Oxaliplatin), PCR outperforms in 3 drugs (Trastuzumab, Temozolomide, and Pemetrexed), TRANSACT outperforms in 2 drugs (Gemcitabine and Paclitaxel), and Combat + Elastic Net outperforms in 1 drug (Cisplatin). No existing method can dominate other methods across different cancer drugs. However, except Cisplatin, the methods (PCR, TRANSACT, and iPCR) that utilize the integrated gene expression data of cell lines and patients through dimension-reduction techniques improve the patient drug response prediction at some levels, compared with Elastic Net and Combat + Elastic Net.

#### Association with patient responses

For assessing the association between our predicted drug responses and actual patient outcomes, we standardized the predicted patient drug-AUC values made by iPCR for each drug (shown in Figure 3). Each pair of violin plots compares these standardized predicted values between responders (red) and non-responders (dark grey). The five pairs of violin plots correspond to the five drugs where iPCR achieves AUC > 0.65. Based on the one-sided Mann-Whitney U test, the significant difference (p-value *<* 0.05) between responders and non-responders in terms of the predicted drug-AUC values by iPCR can be found in Trastuzumab (p-value = 0.0083), Etoposide (p-value = 0.015) and Cisplatin (p-value = 4*×* 10^*−*5^). In addition, the non-responders always have a higher median of predicted drug-AUC by iPCR than the responders for the drugs shown in Figure 3, which is consistent with the fact that a larger drug-AUC is associated with a higher level of drug resistance.

**Fig. 3.**
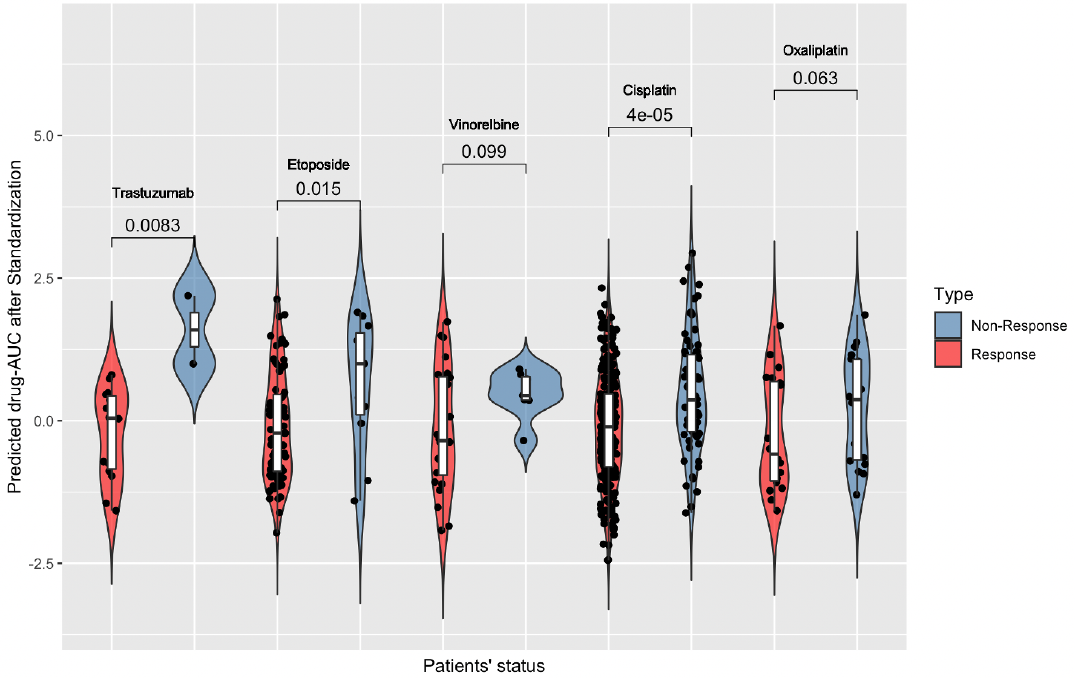
Prediction results of iPCR for five drugs. **A)** For each drug, a pair of violin plots compares the standardized values of predicted patient drug-AUC by iPCR between responders (red) and non-responders (dark gray). The block dots represent the predicted values. The number above each pair of violin plots corresponds to the p-value from the one-sided Mann-Whitney U test.

### 4.3 Shared variation and biological interpretations

In iPCR, given the rank set selected from Algorithm 1, the patient-cell line integration model (Section 2.2.1) helps quantify and compare the amount of shared and individual variation in the gene expression data between cell lines and patients. As shown in Table S1, the patient response prediction for the drugs in Figure 3 uses the rank set 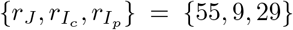 . Given this rank set, the percentage of variation (sum of squares) explained by the estimated joint structure, individual structure and residual noise for cell lines and patients’ gene expression data is shown in Figure 4A. The joint structure is responsible for 59% of variation in both data, which quantify the level of similarities between the genomic profiles of the two biological model systems. The patient gene expression data have a substantial amount of structured variation (18%) that is unrelated to the cell lines, while cell lines’ data only have 5% structured variation that is not shared with patients. This result is consistent with current biological understanding, as the *in vivo* environment is more complex than the *in vitro* environment [61] and various factors that govern it.

In addition, we made use of the biological interpretability of iPCR’s predictors for model validation. Our iPCR model allows an estimation of each gene’s effect on the drug response and hence delineate the underlying biological mechanisms. Based on the ranked list of estimated gene effects, we conducted a PreRanked gene set enrichment analysis [42] for each of the five drugs where iPCR achieves AUC > 0.65, as shown in Figure 4B. We considered two collections of gene sets: 1) Hallmark and 2) Chemical and Genetic Perturbations (CGP). In Figure 4B, each dot corresponds to a gene set, and those gene sets above the dashed line are considered significant with The False Discovery Rate (FDR) *<* 0.05. Based on these significant gene sets identified by iPCR, we observed known drug response mechanisms determined by the expression levels of these genes.

**Fig. 4.**
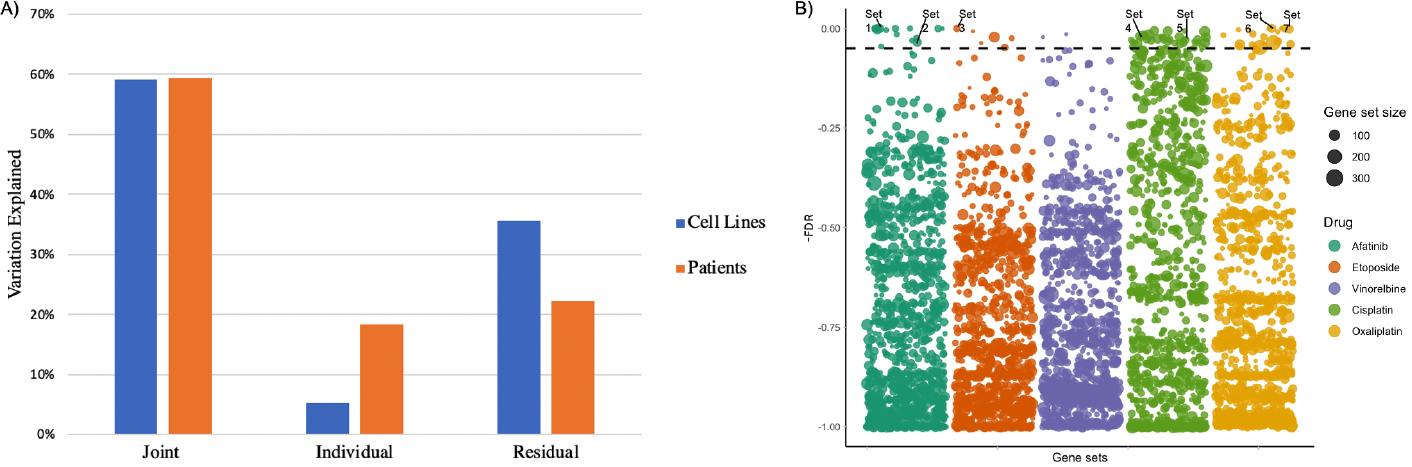
Biological interpretations of iPCR’s results. **A)** Percentage of variation (sum of squares) explained by estimated joint structure (rank = 55), individual structure (rank = 9 for cell lines and rank = 29 for patients) and residual noise for cell lines and patients’ gene expression data. **B)** Bubble plot shows –FDR for PreRanked gene set enrichment analysis (y-axis), based on the estimated gene effects on the patient responses to each of five drugs (in different colors). Each dot corresponds to a gene set, and those gene sets above the dashed line are considered significant (FDR *<* 0.05). The dot size represents the number of genes contained in a gene set. The significant gene sets that are discussed in Section 4.3 are highlighted. Note that Afatinib (for model training) is used as the matching drug of Trastuzumab.

To this end, we investigate seven relevant gene sets from our enrichment analysis that are highlighted in Figure 4B. First, the enrichment of two breast cancer-related gene sets, “Charafe Breast Cancer Luminal vs Mesenchymal Up” [62] (Set 1 in Figure 4B) and “Creighton Endocrine Therapy Resistance 5” [63] (Set 2) has a significant positive association with the sensitivity of Afatinib (FDR *<* 0.001 for Set 1; FDR = 0.034 for Set 2). Set 1 contains HER2 and HER3, and Set 2 contains HER2 and HER4. This finding is consistent with the known mechanism of Afatinib since these three genes are the main targets of Afatinib [64]. Second, the overexpression of genes in the set “Cairo Hepatoblastoma Up” [65] (Set 3) is significantly and positively associated with the sensitivity of Etoposide (FDR *<* 0.001). Set 3 includes the genes upregulated in hepatoblastoma, a cancer type that is typified by the upregulated MYC signaling. Third, the MYC-related gene sets, “Hallmark MYC Targets V1” [66] (Set 4) and “Dang Regulated By MYC Up” [67] (Set 5) play a significant role in the sensitivity of Cisplatin with a positive direction (FDR = 0.031 for Set 4; FDR = 0.024 for Set 5). Our second and third findings are consistent with the past experimental results that Etoposide and Cisplatin provoked a strong apoptotic response in MYC-overexpressing cells [68]. Finally, the p53-related gene sets “Fischer Direct P53 Targets Meta Analysis” [69] (Set 6), and “Hallmark P53 Pathway” [66] (Set 7) are significantly and positively associated with the sensitivity of oxaliplatin (FDR *<* 0.001 for Set 6; FDR = 0.0016 for Set 7); this corresponds to the past finding that colon cancer cells with a wild-type p53 exhibited a characteristic of being sensitive to Oxaliplatin [70].

### 4.4 Gene co-expression network analyses

Using the iPCR results given the rank set 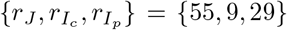 (Table S1), we constructed the shared gene co-expression networks (shown in Figure 5A) as well as cell line-specific and patient-specific networks (shown in Figures S2 and S3 respectively). Furthermore, the genes were clustered into six communities with different colors, based on the Louvain method [71]. We observed two highly dense communities colored in purple and green in the shared network. For downstream interpretations, we considered “hub” genes that significantly connect to more than 30 other genes. We summarize the gene connectivity levels, using a circular heatmap (Figure 5B) that visualizes a matrix with each entry representing the number of edges linked to a hub gene in the shared, cell line-specific or patient-specific network. The dendrogram of genes inside the heatmap shows that the connectivity pattern of genes differs across the shared and model-specific networks, and the shared network contains more hub genes with a high connectivity level (colored in red). The upset plot in Figure S4 shows the number of hub genes in each type of networks as well as their intersections [72]. The names of all hub genes are listed in Table S2.

**Fig. 5.**
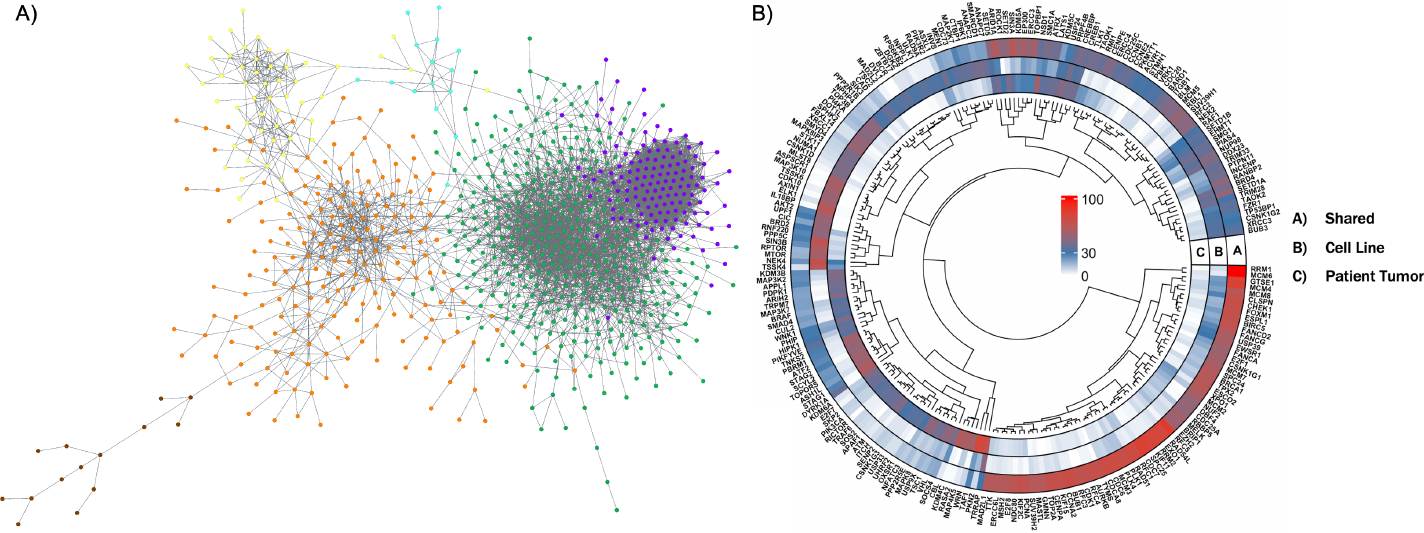
Analysis of model-specific and shared gene co-expression networks between cell lines and patients. **A)** Shared network between the two biological model systems, where each node represents a gene. Significantly associated genes are connected by edges. The gene communities in different colors are detected by an unsupervised clustering method. **B)** Circular heatmap of connection level of each gene in the shared, patient-specific, and cell line-specific networks. Color corresponds to the gene connection level, which is the number of edges linked to a gene (red, high; blue, medium; white low). Only the genes with a connection level > 30 in any of the networks are shown. The dendrogram of genes is provided inside the heatmap.

From our analyses, the top five hub genes in the shared network include RRM1, MCM6, RFC5, SPC25, and EXO1. The hub gene RRM1 provides the precursors that are essential for DNA synthesis, and is functionally associated with DNA replication and DNA damage repair [73]. MCM6 encodes a part of the MCM complex, which is a set of proteins functioning as a helicase [74]. Helicases are essential during DNA replication since they unwind the DNA so that it can be copied. RFC5 relates to homologous DNA pairing and strand exchange, which play a crucial role in repairing DNA damage during mitosis [75]. The upregulation of SPC25 can lead to genetic instability and potentially result in tumorigenesis; its overexpression has been observed in lung cancer, prostate cancer, colorectal cancer and gastric cancer [76, 77]. EXO1 is involved in a number of DNA repair and metabolic pathways [78, 79]. Hence, our estimated shared gene co-expression network between cell lines and patient tumors illustrates that DNA replication and repair are the two critical processes that relate to the pathogenesis of cancer.

## 5 Discussion

We present an integrative multi-system prediction model, iPCR, which serves four main objectives: 1) quantify structured similarities and differences in gene expression data between cell lines and patient tumors, 2) exploit patient-cell line similarities to predict patient drug response, 3) identify key genomic drivers for drug-specific patient response, and 4) infer both model-specific (cell lines or patients) and shared gene co-expression networks while detecting hub genes within these networks. We demonstrate that accounting for model-specific information in the prediction model not only enhances the accuracy of drug response prediction in both simulation studies and real-world applications, but also improve the interpretability of shared biology and disparities between cancer model systems.

Our iPCR model can be generalized in several directions. First, our framework can be potentially used to integrate multiple cancer model systems—such as cell lines, patient-derived xenografts (PDXs), and patient tumors—into a single statistical framework. This enables the investigation of their common biology and disparities while predicting patient drug responses. PDXs, derived from human tumors and cultivated in mice, are emerging as systems that more faithfully represent human tumors [80]. However, there are still biological differences between PDXs and patient tumors, as PDXs may not consistently capture the dynamic shifts in copy number alterations found in primary human tumors. [80]. One could reformulate the iPCR model based on Structural Learning and Integrative Decomposition of Multi-View Data (SLIDE) [39] to study the shared, partial-shared, and individual features among the three different but related cancer model systems. Second, our iPCR model could potentially be improved to allow out-of-sample prediction. In the current framework, although patient drug response data are not utilized in model training, our model only predicts drug response for patients whose genomic data are used for model training. For a new patient, the joint score can be obtained by projecting the patient’s gene expression data onto the shared loading, which can then be utilized for predicting the drug response. However, both the quantification and preprocessing of the new patients’ genomic data require careful consideration. Finally, our shared gene co-expression network is constructed based on two joint sample covariance matrices. It would be of interest to refine the current sample covariance decomposition method to obtain a single joint sample covariance matrix for better interpretation. To achieve this goal, additional distributional assumptions might be required.

The iPCR package is available at https://github.com/bayesrx/iPCR. Guidelines for downloading the original data, as well as the preprocessed data, are also provided through this GitHub link.

## 6 Acknoledgement

This work was partially supported by R01HG010731 (GL), P30 CA046592 (GL, VB) and R01-CA244845-01A1 (VB).

## Supplementary Materials

In the supplementary materials, we first address the estimation of the gene co-expression network in Section S1, including the proof of the uniqueness and identifiability for the covariance decomposition. Following this, Section S2 provides additional details on our simulation studies, specifically focusing on the sensitivity analysis of rank selection for iPCR. Lastly, Section S3 showcases additional results from our pan-cancer application.

### S1 Gene co-expression network estimation

#### S1.1 The solution of sample covariance decomposition

For all *i ∈*{*c, p*}, the unique solutions of 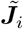 and 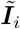 that meet all the conditions in Proposition 2 are given by

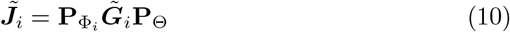

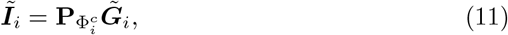

where 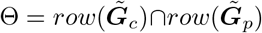 is the shared feature space, 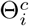 is the orthogonal complement of Θ in *row*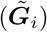, Φ_*i*_ is the orthogonal complement of *col*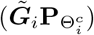 in *col*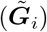, and 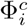 is equal to 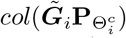. Based on the proof of Proposition 1 in [40], Θ is equivalent with *row*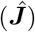, and 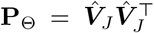. The projection matrices 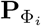and 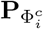 can be easily acquired following their definitions.

Note that each column of 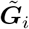(a gene feature) always has a mean of 0 due to the centralization for each column of **C** and **P** as well as the JIVE estimation algorithm [38] in iPCR Stage I. Then, based on Corollary 1 in Section S1.2, the unique solutions for 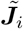 and 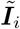satisfy the property of sample covariance decomposition as described in Model 6.

#### S1.2 Uniqueness and identifiability of the decompostion solution

##### Lemma 1

*Given a set of matrices {A, B,C}, if A = B + C and rank(A) = rank(B) +rank(C), then row(A) = row(B) ∪ row(C) and col(A) = col(B) ∪ col(C).*

*Proof* : Denote *a = rank(A), b = rank(B), c = rank(C), and dim(A)* as *m*×n. We first prove *row(A) = row(B) ∪ row(C)* by showing *row(A) ⊆ row(B) ∪ row(C)*, and *row(B) ∪ row(C) ⊆ row(A)*.

1. It is straightforward to show that *row(A) ⊆ row(B) ∪ row(C)* since each row of *A* is a linear combination of the corresponding rows of *B* and *C*.
2. To show *row(B) ∪ row(C) ⊆ row(A)*, first do linear transformation of rows in *B* to get *B*^*′*^ where only first *b* rows are nonzero, such that 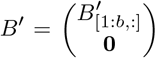. Then do linear transformation of rows in *C* to get *C*^*′*^ such that only *c* rows are nonzero, and the nonzero rows in *B*^*′*^ and *C*^*′*^ do not have a common index (because of the rank constraint), such that 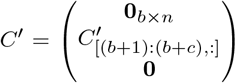 . Those c rows, which form the basis for *row(C)*, belongs to *row(A)* since the corresponding rows in *B*^*′*^ has been zeroed out. Similarly, it can be easily shown that *row(B)* also belongs to *row(A)*. Hence, *row(B) ∪ row(C) ⊆ row(A)*.

Same proof can be used to show *col(A) = col(B) ⋃ col(C)* after transpose *A, B*, and *C*. □

##### Proposition 2

Given a set of matrices 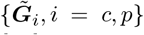, there are unique sets of matrices 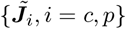 and 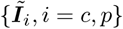 such that,

1. 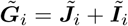 for i = c, p
2. 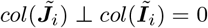 for i = c, p
3. 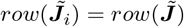 for i = c, p, where 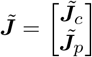
4. 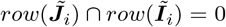 for i = c, p
5. 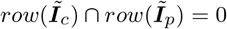

*Proof* : Let *i = c, p* for this proof. Define 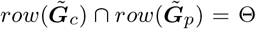, and 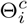as the orthogonal complement of Θ in 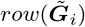.

##### Existence

When 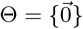:

It is obvious that Condition 1-5 are all satisfied as 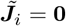, and 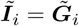.

When 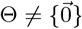:

Define Φ_*i*_ as the orthogonal complement of *col*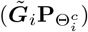 in *col*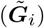, and 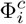as *col*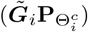. We can derive that

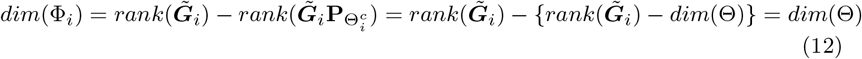

Then we will show that 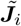and 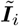defined below satisfy Condition 1-5:

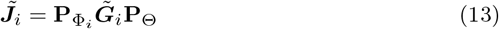

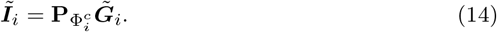

Condition 1:

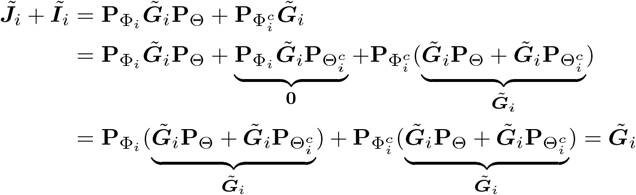

Condition 2: Since Φ_*i*_ is orthogonal with 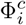, then Condition 2 is satisfied.

Condition 3:

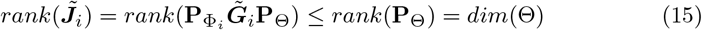

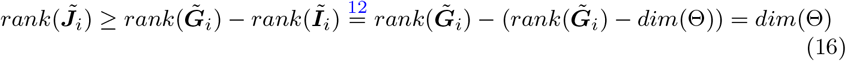

Based on Equation 15 and 16, we can get 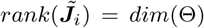. Since we also know that 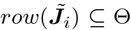 from Equation 13, we can conclude that 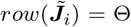 and 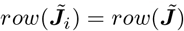.

Condition 4: Since 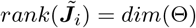and 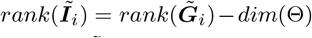, then

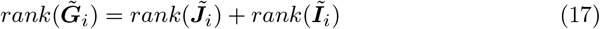

Thus, Condition 4 is satisfied based on the theorem from [81].

Condition 5: Since 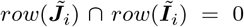from Condition 4 and 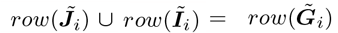 from Lemma 1, then 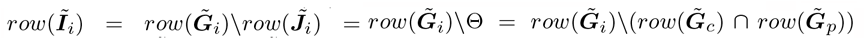. Hence, 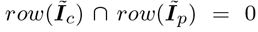.

##### Uniqueness

Assume there exist another 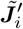 and 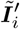 satisfying Condition 1-5 such that 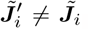, 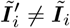 from part 1. Since 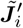 and 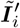 satisfy Condition 1, 2 and 4, then

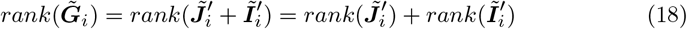

based on the theorem from [81].

Based on Equation 18, Lemma 1 and Condition 3-4, we have

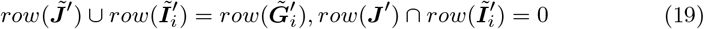

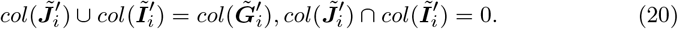

By integrating Equation 19 with Condition 5, we can easily show 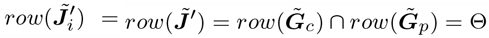.

Based on Equation 20 and Condition 2, we can rewrite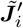 and 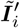 as

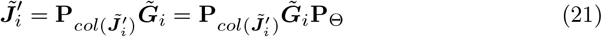

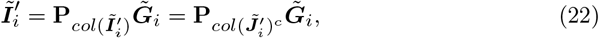

where 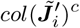 is the orthogonal complement of 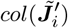 in 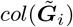.

By combining Equation 21, 22 and Condition 1, we have

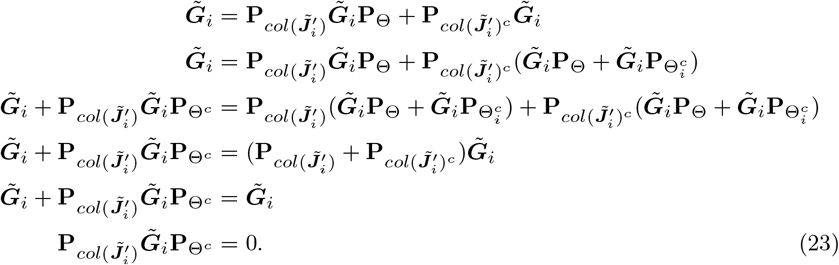

Since 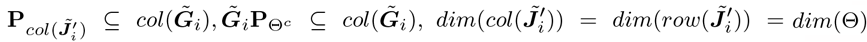, and 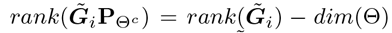, then 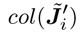 must be equal to the orthogonal complement of 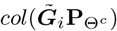 in 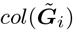, which means 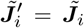 and 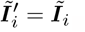. □

###### Corollary 1

*Suppose a set of matrices* 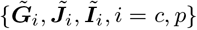 *that meets all the conditions in Proposition 2. For all* i *∈ {*c, p*}, if* 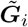*is column-centered by mean within each column (feature), and samples in* ***G****i are assumed to be independent and identically distributed, then* 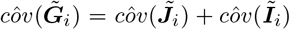 *with respect to the sample covariance matrix between features*.

*Proof* : Based on the proof of Proposition 2, we know 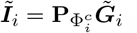.

Since 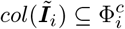 and 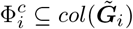, then 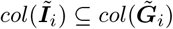.

Since the linear combination of centered columns is still column-centered, ***Ĩ***_*i*_ is column-centered. Then, 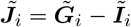 is also column-centered.

Define 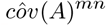 and 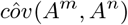 as the sample covariance between *m*^*th*^ feature and *n*^*th*^ feature in matrix A. The covariance equation below is always satisfied:

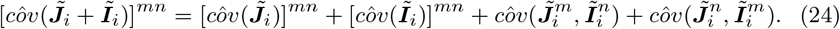

Based on Condition 4 in Proposition 2, and the column-centered 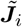 and 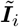, we have

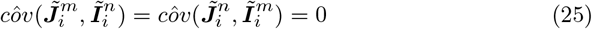

Hence, 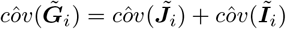. □

### S2 Additional information of simulation studies

#### S2.1 Data generating process

To construct the SVD components of ***J***_*c*_, ***J***_*p*_, ***I***_*c*_, and ***I***_*p*_ in Model 7 and 8, we first generate elements of 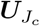 and 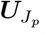 from *N* (0, 4), and elements of ***V***_*J*_ and ***V***_*J*_ from *N* (0, 1). We subsequently center and orthonormalized 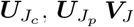 and ***V***_*J*_ . Then, we set **Σ**_*J*_ = *diag*(40, …, 10), where the 40 diagonal elements in **Σ**_*J*_ are equally spaced. Next, we generate elements of 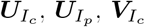 and 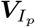from *N* (0, 1), and subsequently center and orthonormalized them, where 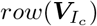 and 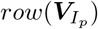 are constrained to be orthogonal with each other as well as with *row*(***V***_*J*_). After that, we set 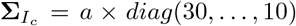 and 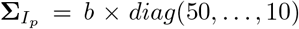, where the 30 diagonal elements in 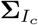and 50 diagonal elements in 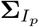 are equally spaced. The values of *a* and *b* can be easily calculated to meet different levels of joint proportion. Constructing in this way, we have 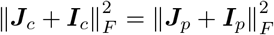, which avoids the issue of “the bigger dataset always wins” during data integration. Last, all these SVD components are scaled down by 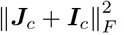 .

#### S2.2 Sensitivity analysis of rank selection for iPCR

To better understand the results of sensitivity analysis in Figure S1, first we note that there are three types of underlying data structures in our simulated genomic data of cell lines and patients, including the joint variation, the model-specific variation, and the residual noise. The joint rank *r*_*J*_ corresponds to *r*_*J*_ latent joint principal components in the joint variation. The individual ranks 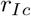 and 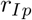 correspond to 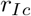 and 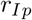 latent individual principal components in the cell line-specific and patient-specific data, respectively. A value of joint proportion that is close to 1 (0) illustrates that each joint principal component explains a higher (lower) proportion of variation in the data than each individual principal component does. In addition, a *SNR*_*stage*1_ that is much less (higher) than 1 demonstrates that each of those top principal components from the noise could explain higher (lower) or comparable proportion of variation in the data than some joint and individual principal components do. We call those top principal components in the matrix of noise as residual principal components.

The sensitivity of rank selection to prediction accuracy is affected by the proportions of variation explained by joint, individual and residual principal components. Let 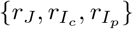 be the form of a rank set. First, when joint proportion is larger than 0.6, the three subplots in Figure S1 show that the sensitivity of rank selection to the prediction accuracy is relative low except {30, 40, 60}. This is expected since in this case the top principal components in the data correspond to the joint principal components. Hence, almost all joint principal components that relate to the drug response are easy to be detected, even though the joint rank is slightly overestimated or the individual ranks are slightly misspecified. The underestimation of joint rank could omit important predictors in the drug response prediction model, and significantly reduce the prediction accuracy. Second, when *SNR*_*stage*1_ = 0.5, the sensitivity of rank selection to the prediction accuracy is relative low except {30, 40, 60} under high joint proportion. One possible reason is that since the noise is large and some residual principal components explain comparable proportion of variation in the data with the joint and individual principal components, only part of the joint principal components could be accurately detected even under the true ranks. Hence, slightly misspecified the ranks do no have large effect on the prediction accuracy. Third, when *SNR*_*stage*1_ increases from 1 to 2, the prediction accuracy of iPCR given {40, 40, 60} (overestimation of individual ranks) decreases under low joint proportion. One possible reason is that when *SNR*_*stage*1_ = 1, the top residual principal components explain comparable proportion of variation in the data with those bottom individual principal components; hence the overestimation of individual ranks treats those residual principal components as individual principal components, and it does not affect the model estimation too much to detect the joint principal components. In contrast, when *SNR*_*stage*1_ = 2, the variation in the noise is relatively low, and the top residual principal components could not be appropriately treated as individual principal components during the model estimation; hence, the overestimation of individual ranks significantly affect the prediction accuracy in this case.

**Fig. S1.**
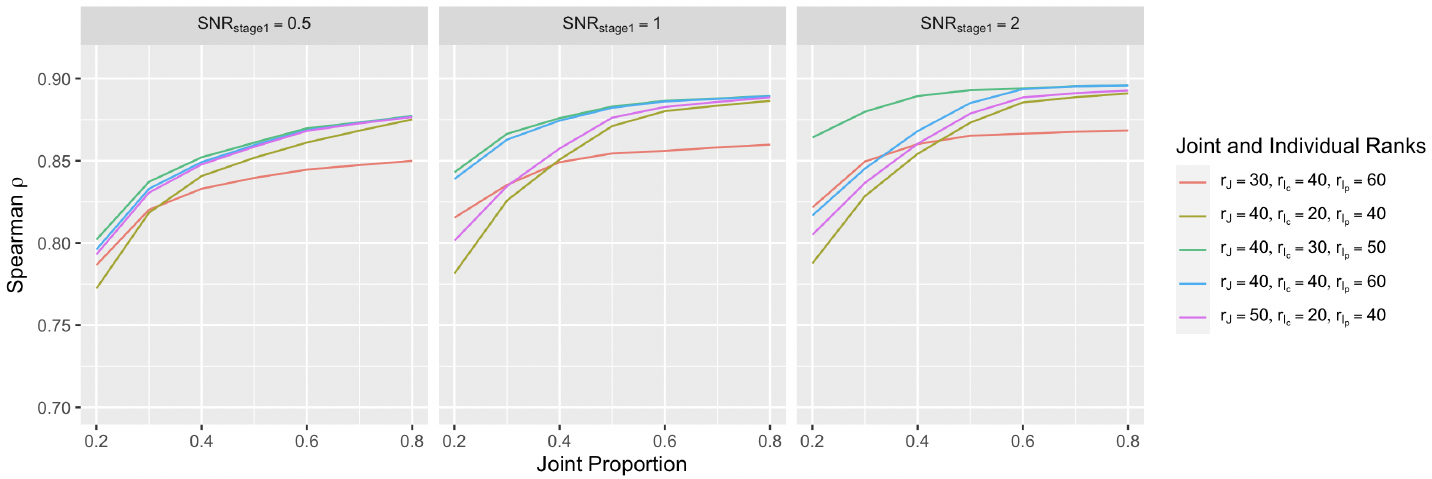
Accuracy comparison of iPCR under different rank combinations in predicting **D**_**p**_ in simulation study. The prediction performance is measured by Spearman correlation between the true and predicted values of **D**_**p**_. The data are generated under different levels of *SNR*_*stage*1_ and joint proportion over 50 replications. The solid curve corresponds to the sample mean of Spearman correlation over replications.

### S3 Additional results of pan-cancer application

#### S3.1 Patient drug response by iPCR

**Table S1.** Rank selection results of iPCR models for drug response prediction in patients.

| Drug | Joint rank | Individual rank of cell line | Individual rank of patient |
| --- | --- | --- | --- |
| Trastuzumab (Afatinib) | 55 | 9 | 29 |
| Etoposide | 55 | 9 | 29 |
| Vinorelbine | 55 | 9 | 29 |
| Cisplatin | 55 | 9 | 29 |
| Oxaliplatin | 55 | 9 | 29 |
| Gemcitabine | 55 | 9 | 29 |
| Temozolomide | 45 | 19 | 39 |
| Pemetrexed | 55 | 9 | 29 |
| Paclitaxel | 45 | 19 | 39 |

*Note:* The drug name in parenthesis corresponds to the matching drug from GDSC. For each drug, we report the joint and individual ranks selected by iPCR model through Algorithm 1 among the rank set candidates mentioned in Section 4.2

#### S3.2 Gene co-expression network estimation

**Fig. S2.**
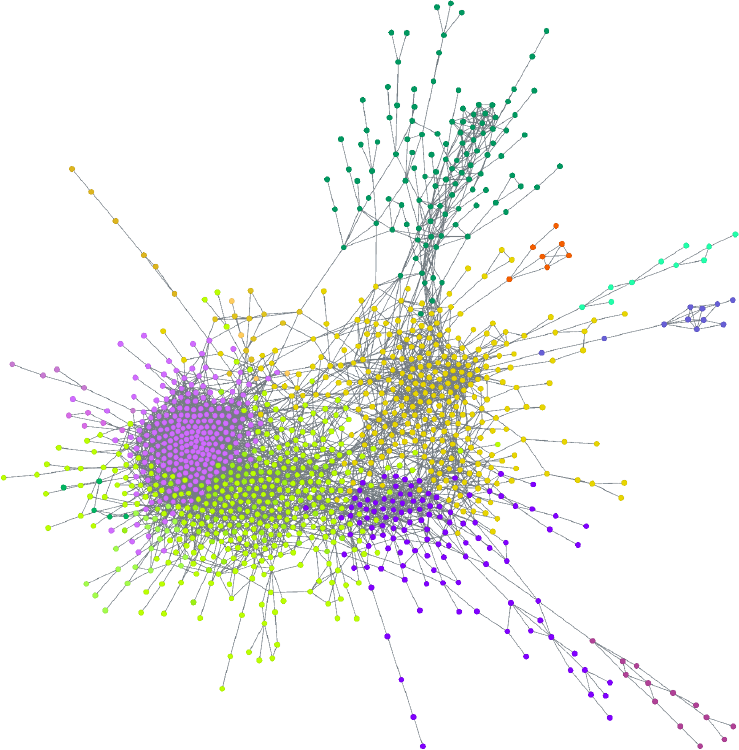
Cell line-specific gene co-expression network, where each node represents a gene, and those significantly associated genes are connected by edges. The gene communities in different colors are detected by an unsupervised clustering method.

**Fig. S3.**
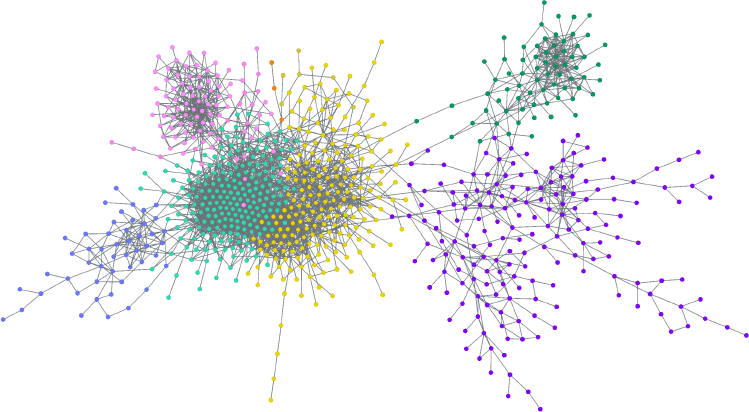
Patient-specific gene co-expression network, where each node represents a gene, and those significantly associated genes are connected by edges. The gene communities in different colors are detected by an unsupervised clustering method.

**Fig. S4.**
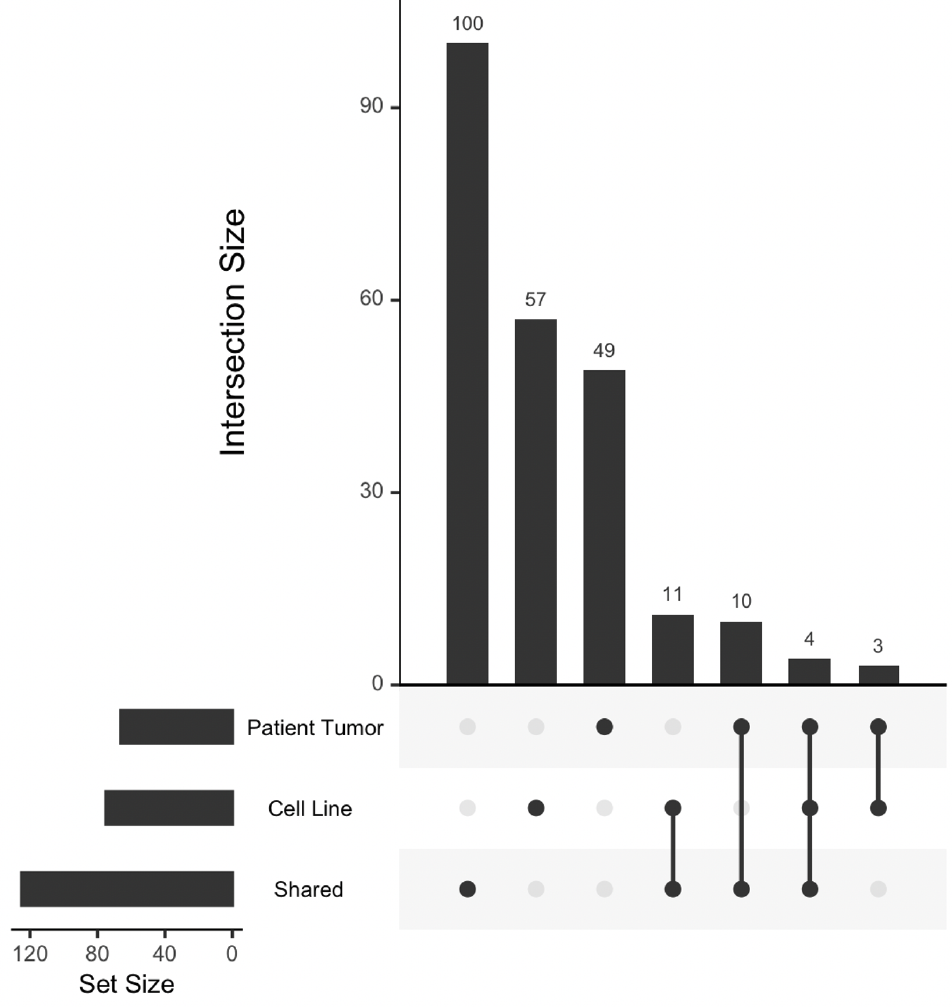
The upset plot lists the number of hub genes (those genes that significantly connect to >30 other genes) in each type of networks and their intersections.

**Table S2.**
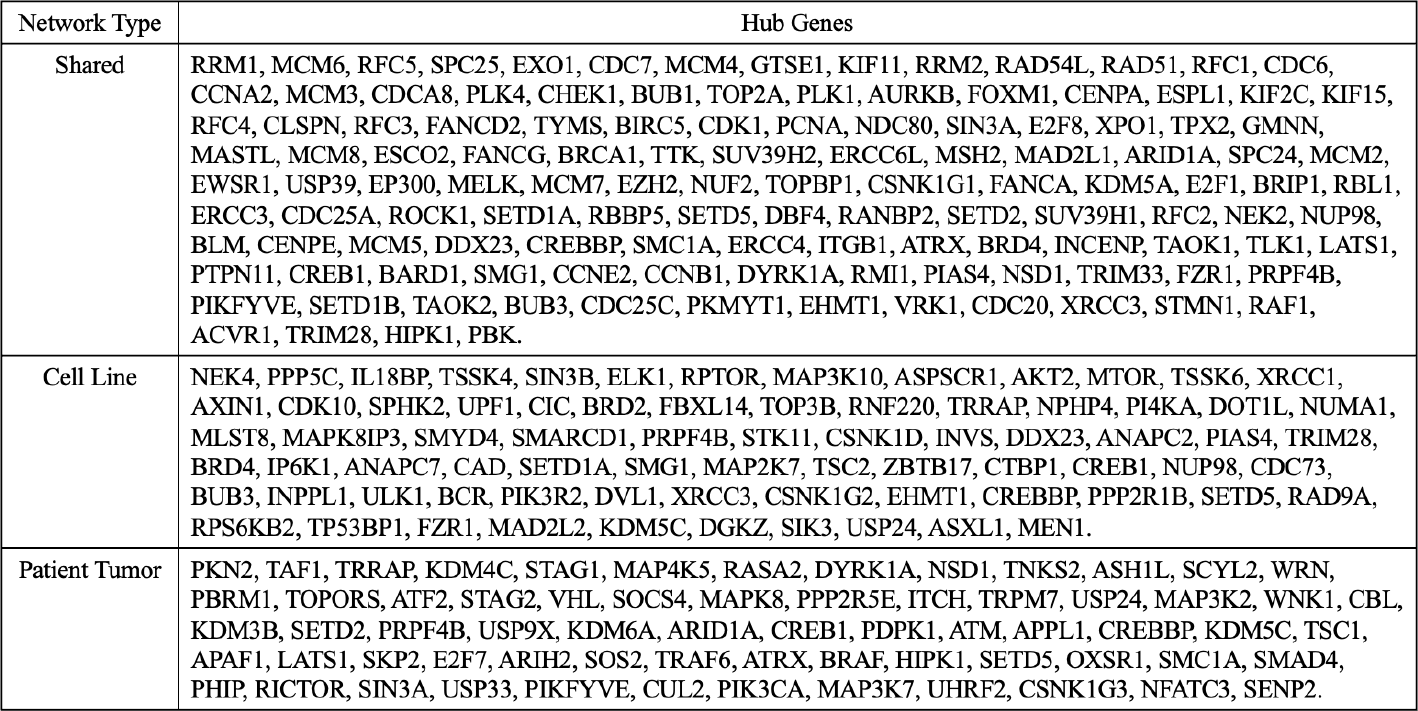
Hub genes in shared, cell line-specific, and patient-specific gene co-expression networks.

## Notes

### Competing Interest Statement

The authors have declared no competing interest.

